# Genome-scale metabolic rewiring to achieve predictable titers rates and yield of a non-native product at scale

**DOI:** 10.1101/2020.02.21.954792

**Authors:** Deepanwita Banerjee, Thomas Eng, Andrew K. Lau, Brenda Wang, Yusuke Sasaki, Robin A. Herbert, Yan Chen, Yuzhong Liu, Jan-Philip Prahl, Vasanth R. Singan, Deepti Tanjore, Christopher J. Petzold, Jay D. Keasling, Aindrila Mukhopadhyay

## Abstract

Achieving high titer rates and yields (TRY) remains a bottleneck in the production of heterologous products through microbial systems, requiring elaborate engineering and many iterations. Reliable scaling of engineered strains is also rarely addressed in the first designs of the engineered strains. Both high TRY and scale are challenging metrics to achieve due to the inherent trade-off between cellular use of carbon towards growth *vs.* target metabolite production. We hypothesized that being able to strongly couple product formation with growth may lead to improvements across both metrics. In this study, we use elementary mode analysis to predict metabolic reactions that could be targeted to couple the production of indigoidine, a sustainable pigment, with the growth of the chosen host, *Pseudomonas putida* KT2440. We then filtered the set of 16 predicted reactions using -omics data. We implemented a total of 14 gene knockdowns using a CRISPRi method optimized for *P. putida* and show that the resulting engineered *P. putida* strain could achieve high TRY. The engineered pairing of product formation with carbon use also shifted production from stationary to exponential phase and the high TRY phenotype was maintained across scale. In one design cycle, we constructed an engineered *P. putida* strain that demonstrates close to 50% maximum theoretical yield (0.33 g indigoidine/g glucose consumed), reaching 25.6 g/L indigoidine and a rate of 0.22g/l/h in exponential phase. These desirable phenotypes were maintained from batch to fed-batch cultivation mode, and from 100ml shake flasks to 250 mL ambr® and 2 L bioreactors.

## Introduction

Heterologous production of bioproducts has been demonstrated for a very large number of compounds and in a wide variety of microbial hosts^1,2^. Yet, even the most well-designed heterologous pathway requires considerable additional work to reach titers, rates and yields (TRY) necessary for the adoption of these systems by industry^3,4^. In addition, the production parameters of a strain at lab-scale is often not predictive of its performance and robustness when cultivated in different modes or at larger scales. As a result, only a small fraction of bioproduction strains have been successfully scaled and deployed^2^.

Here we explore if it is possible to rewire the metabolism of the host strain such that production of a final product or a key intermediate is coupled with the carbon source, and used to maximize and maintain productivity at scale. Native microbial processes that take such growth coupled routes include the generation of ethanol and organic acids during fermentation. Production of these metabolites are required for carbon utilization during fermentative growth, and correspondingly these compounds represent the most prominent examples of successful high-volume bioproduction^5,6^. We hypothesized that coupling production to growth is implementable for a heterologous product, and that such a dependence could provide high TRY and the ability to maintain production parameters across different growth modes and scales.

The availability of comprehensive metabolic models and genome editing tools in a wide variety of microbes suitable for industrial use provides the foundation for our approach. We use the production of indigoidine, a bipyridyl compound derived from glutamine, as the target heterologous product. Both as a sustainable replacement for blue pigments^7^ in a wide array of applications as well as a model non-ribosomal peptide^8^, this compound provides a valuable target to explore. We used *Pseudomonas putida* KT2440 as our production host, leveraging the availability of the iJN1462 genome scale model for *P. putida* KT2440^9^. We adapted elementary mode analysis (EMA)^10^ to determine the constrained minimal cut set (cMCS) required to minimize metabolic flux towards undesired products and link indigoidine formation to cell viability^11^. These analyses, combined with publicly available omics data^12,13^, provided the set of gene loci that represented the reactions necessary for removal. The corresponding set of gene loci were repressed using multiplex CRISPR interference (CRISPRi) that we optimized for use in *P. putida* KT2440. Our implementation resulted in a highly edited strain that, in a single iteration of strain engineering, achieved close to 50% max theoretical yield of indigoidine in *P. putida* KT2440 and TRY characteristics that maintain fidelity from laboratory to industrially relevant scales.

## Results

### Genome scale evaluation of P. putida for strong coupling

To develop the product coupling approach (**Figure 1a**), we first identified the number of metabolites represented in *P. putida* iJN1462^9^ model that can be made essential for growth. For this we used the cMCS algorithm^11^ that can identify minimal sets of reactions, the elimination of which would cause production of a given metabolite to become essential for growth. Aerobic conditions with glucose as the sole carbon source were used to model growth parameters. We searched for gene knockdown sets to satisfy three potential constraints in which the theoretical product yield was at a minimum of 10%, 50%, or 80% of the maximum theoretical yield (MTY) for all producible metabolites in the model coupled to a minimum 10% biomass yield. This analysis, completed for all 2145 metabolites in the genome scale model, indicated that 979 organic metabolites could potentially be made essential for growth. In the first pass, 98.6% of these 979 metabolites had the potential to satisfy this biomass-formation constraint, with a minimum production threshold of 10% MTY. When the threshold for minimum production was set to 50% MTY, 903 metabolites could be essential for growth; for an 80% threshold MTY, only 444 metabolites could be made essential for growth, representing only 45% of the total producible metabolites. This potential growth coupling for all metabolites is consistent for similar calculation for other hosts^11^ (see **Supplementary Table 1)**. Setting a higher demand for minimum product yield results in fewer metabolites that can be used to implement a production obligatory regime.

**Table 1:** Maximum theoretical yield (MTY) of glutamine and indigoidine from two different substrates glucose and galactose with respect to stoichiometry and redox balance in *P. putida*

| Metabolite | Maximum theoretical Yield (MTY) |  |  |  |
| --- | --- | --- | --- | --- |
|  | Glucose |  | Galactose |  |
|  | mol product/<br>mol substrate | g product/<br>g substrate | mol product/<br>mol substrate | g product/<br>g substrate |
| Alpha-ketoglutarate | 1.320 | 1.07 | 1.366 | 1.11 |
| Glutamine | 1.141 | 0.93 | 1.181 | 0.96 |
| Indigoidine | 0.537 | 0.74 | 0.556 | 0.77 |

**Figure 1:**
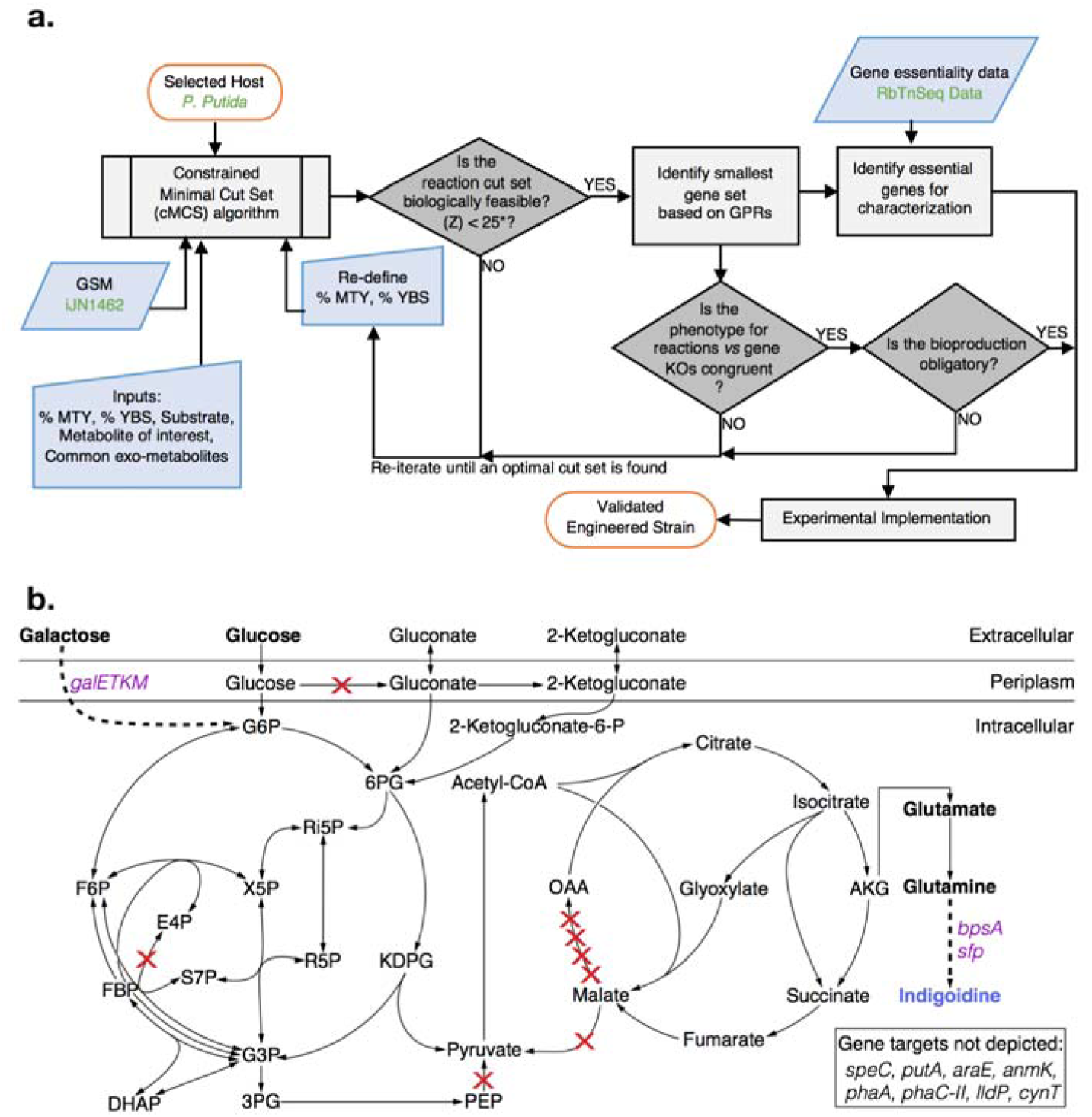
Computationally Guided Predictions for Metabolic Rewiring in *P. putida*. **a**, Modeling and engineering workflow diagram. This approach can potentially be extended to any carbon source, host and/or metabolite. Input specific to this specific host/final product work is marked in green font. **b**, The central metabolism of *Pseudomonas putida* engineered to produce indigoidine from either glucose or galactose. Heterologous genes are indicated in purple text. Indigoidine is derived from the TCA intermediate alpha-ketoglutarate (AKG) via two molecules of glutamine. The genes targeted in *P. putida* central metabolism for knockdown by dCpf1/CRISPRi are indicated with red X marks. Additional gene targets outside of *P. putida* central metabolism are indicated in the box on the bottom right. A total of 14 genes were targeted for CRISPR interference excluding *mqo-I* and *cynT*, as the latter are essential by genome-wide transposon mutagenesis (RB-TnSeq). Refer to **Supplementary Table 4** for more information.

As the framework proposed by von Kamp and Klamt required an extensive rewiring of microbial physiology, *a priori* it was not obvious how to account for a heterologous end product, a challenge to implement in practice. We began by adding an *in silico* reaction for the heterologous product, indigoidine, to the genome scale metabolic model iJN1462^9^. This reaction represents the biosynthesis of indigoidine from glutamine and accounts for all necessary cofactors. The MTY for glutamine and indigoidine was calculated to be 1.141 mol/mol and 0.537 mol/mol respectively from glucose as the carbon source (**Table 1**). The MTY for glutamine in *P. putida* was high relative to other hosts screened by us (**Supplementary Table 2**). As this method accounts for the other physiological processes competing for resources, a MTY derived from a genome scale model provided a more accurate assessment compared to simpler methods, as is commonly done in the field^14,15^.

In order to predict reactions that would be required to improve indigoidine production, we used glutamine, its precursor, to conduct the analysis. Our process for determining the list of required gene targets is diagrammed in **Figure 1**. The minimum theoretical product yield of glutamine was set at 10%, 50% and 80% MTY to derive the reactions that would require knockout or knockdown for product substrate paired growth. We eliminated potential target sets that needed the removal of genes coding for multi-functional proteins, as we sought to limit additional metabolic perturbations that could confound our analysis. Of the 1956 reactions in iJN1462 that are associated with genes, only 60% have a single gene associated with them. If a metabolic reaction was catalyzed by more than one gene product (genes coding for isozymes or multi-subunit enzymes), we included both genes for inactivation. After implementing these filters, we found that a threshold of 80% MTY could be achieved using the elimination of 14 cellular reactions. These 14 metabolic reactions when mapped to their corresponding genes and gene products represented 16 gene loci (**Figure 1b and Supplementary Dataset 1**).

Next, we used Flux Balance Analysis (FBA) and Flux Variability Analysis (FVA), to confirm the 16-gene cMCS strategy to be obligatory for glutamine production. Using our constructed cMCS platform, we set the parameters to explore potential product-obligatory strategies to enhance the production of indigoidine in *P. putida* when glucose is fed as the sole carbon source. This 16 gene set provided for glutamine was then extended to assess production paired growth for indigoidine. FBA analyses confirmed that growth using glucose could be paired with indigoidine production at 90% theoretical yield (0.48 mol/mol or 0.66 g/g of glucose). This analysis also confirmed that we can adapt the work from von Kamp and Klamt^11^ for non-native final products and target specific genes rather than enzymatic reactions for intervention.

Since EMA requires the delineation of specific growth conditions, such as starting carbon source, we examined if the gene cut set with glucose as a substrate, could maintain product pairing with other known native carbon substrates for *P. putida*, such as *para*-coumarate and lysine^12,16^. FBA with these alternate carbon sources (i.e. lysine, *para-*coumarate) indicated that a strain engineered using the 16-gene cMCS strategy for the glucose would fail to produce glutamine (**Supplementary Table 3**). In contrast, this gene targeting set (**Supplementary Dataset 1**) results in the desired production obligatory growth using galactose as a carbon source because it shares the same downstream catabolism as glucose (**Figure 1b**).

### Building the multi-edit engineered strain

To test the predictions from the metabolic modeling described above, we built the control engineered *P. putida* production systems. First we genomically integrated the heterologous production pathway comprised of *sfp* and *bpsA*. BpsA is a non-ribosomal peptide synthetase (NRPS) from *Streptomyces lavendulae* that catalyzes indigoidine formation from two molecules of glutamine in an ATP-dependent manner^17^. Activation of BpsA requires a post-translational pantetheinylation conferred by a promiscuous Sfp from *Bacillus subtilis*^*18*^. The genomically integrated production strain harboring a plasmid-borne *dCpf1* and non-targeting gRNA serves as the control production strain. The basal production of indigoidine in *P. putida* is 2.3 g/L indigoidine from 10 g/L glucose after 24 hours (**Supplementary Figure 1a**). The bulk of production occurred in stationary phase, approximately 12 hours after carbon depletion, as is typically observed for *P. putida*^*19,20*^. To test the use of galactose, we also engineered a galactose utilization strain via genomic integration of a *galETKM* operon^21,22^ and here production of indigoidine was negligible (**Supplementary Figure 1b**). Optimizing carbon/nitrogen ratio yielded only modest improvements to indigoidine production for both glucose and ammonium sulfate (**Supplementary Figure 1c-e**).

Prior to construction of the multi-gene edited production strain, we assessed if our gene set contained essential genes. The iJN1462 model has an incomplete list of essential genes; in addition we manually annotated genes as essential or dispensable using gene essentiality data generated from a barcoded fitness library (RB-TnSeq)^13^ **(Supplementary Dataset 2)**. Out of the 16 genes identified for knockdown, two genes were excluded because they are essential for viability. By eliminating essential genes from the targeted gene set, we hypothesized that the predicted metabolic rewiring is more consistent with product substrate pairing rather than growth coupling.

To efficiently overcome technical limitations required to make 14 gene edits, we implemented a multiplex CRISPRi/dCpf1 targeting strategy. We drew on our understanding of repetitive element instability^23,24^ to minimize use of repeated DNA sequences to limit gRNA array loss. While other reports have indicated technical challenges constructing multiplex gRNA arrays^25^, both native and synthetic repetitive arrays exist (including those of native CRISPR arrays)^26,27^. An endonuclease deficient class II CRISPR-Cas enzyme, FnCpf1^28^, was chosen over Cas9 as the Cpf1 crRNA is only 19 bp in size, compared to the corresponding crRNA (gRNA scaffold sequence) from Cas9, which is 76 bp^28^. Each gRNA was driven by a different *P. putida* tRNA ligase promoter/terminator pair, and dCpf1 was placed under the control of the *lacUV5* promoter. Minimal 100-bp promoter sequences from native tRNA ligases were sufficient to express *mCherry* fluorescent protein, confirming that heterologous mRNA transcripts for gRNAs would be generated (**Supplementary Figure 1f**).

In a successful deployment of the multiplex CRISPRi/dCpf1 we expect to see a decrease in mRNA expression levels (and protein abundance) of the genes targeted with CRISPR interference. We used RNAseq analysis to examine the engineered strain, and compared normalized RNA expression levels between the control strain (**Figure 2a-c**). RNA expression levels were sampled over the duration of a 72-hour time course. Expression of all 14 gRNAs were detected by this analysis (**Figure 2a**). The multiplexed Cpf1 gRNAs in this array did not efficiently terminate with endogenous terminator sequences and generated chimeric mRNAs. Nonetheless, nine of the fourteen targeted gene loci exhibited decreased mRNA expression levels, and at best showed a 50% decrease **(Figure 2b, Supplementary Figure 2)**. Global indirect changes in gene expression were also detected (**Figure 2c**). Partial reduction of protein abundance was also observed for ten of the fourteen genes **(Figure 2b, Supplementary Figure 2**).

**Figure 2:**
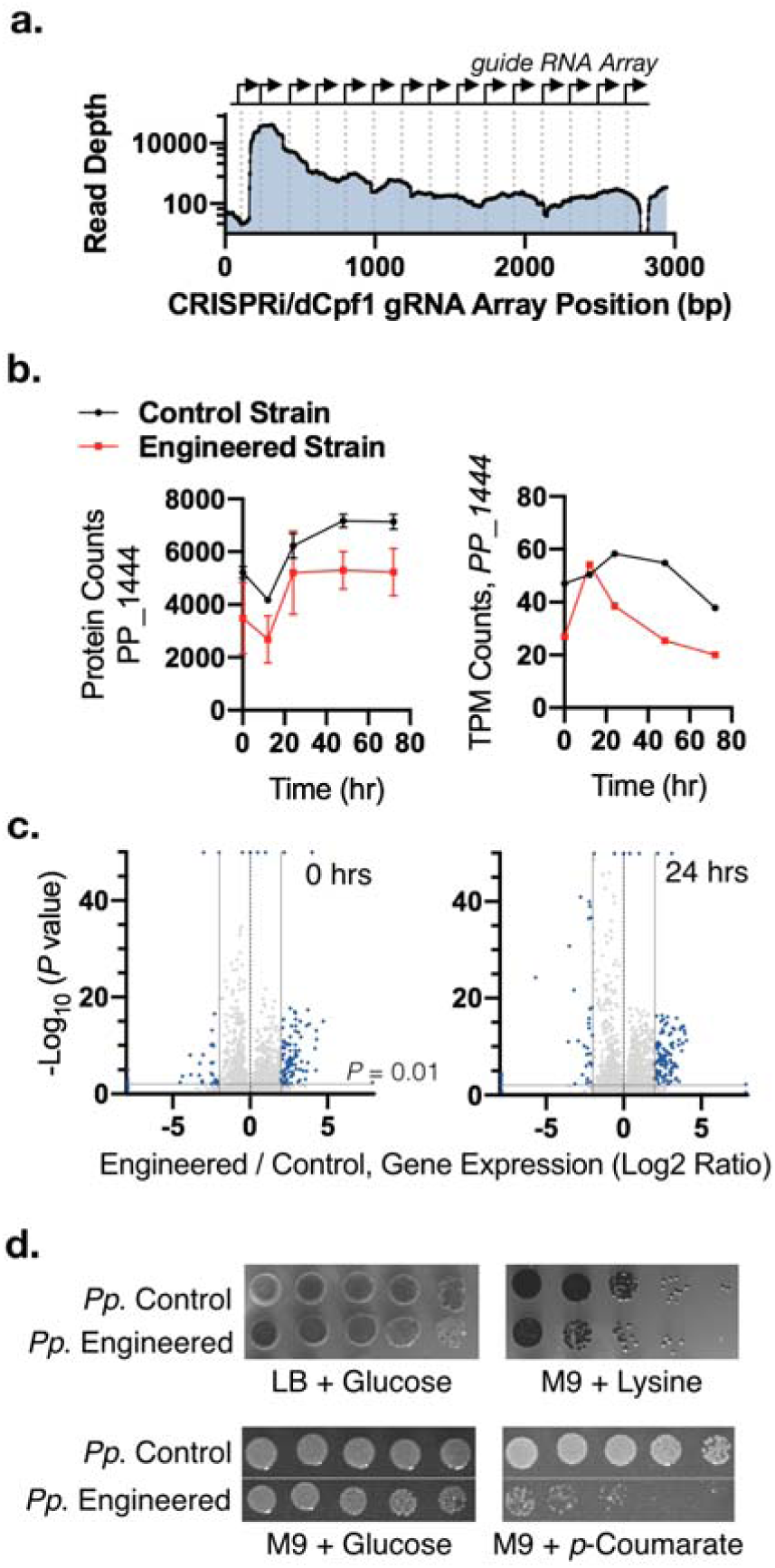
Characterization of the multi-gene engineered strain via RNAseq and Proteomics. **a-c**, *P. putida* harboring a genomically integrated indigoidine expression cassette and either an empty vector (control strain) or a dCpf1/CRISPRi targeting array examined for gene knockdown efficiency. **a**, RNAseq analysis of plasmid-borne gRNA array in *P. putida*. **b**, Knockdown efficiency of a representative gene locus targeted for inhibition over a 72-hour time course. RNA expression levels (right hand panel) were validated with targeted proteomic analysis (left hand panel). Proteomic samples were analyzed in at least biological triplicate. RNAseq analysis for the control sample was completed in biological duplicate for the control and biological quadruplicate for the engineered strain. **c**, dCpf1/CRISPR interference causes global RNA expression level changes. Volcano plot of mRNA expression levels compared at t = 0 h and t = 24 h between multi-gene engineered and control strains. 184 datapoints (0 hr) and 391 datapoints (24 hrs) out of 5369 datapoints are outliers and are some are displayed on the edge of the axes. **d**, Validation of carbon source rewiring. Genome-scale modeling predicts that glucose/indigoidine rewiring blocks growth of engineered strains on lysine as a carbon source.

A consequence of pairing production to the catabolism of specific carbon sources, results in predictions that other carbon sources can no longer be metabolized **(Supplementary Table 3)**. We tested this prediction experimentally and observed that engineered strains for product substrate pairing showed reduced growth when using either lysine or *para-*coumarate as the sole carbon source, in agreement with the modeling (**Figure 2d**).

### Characterizing the multi-gene engineered production strain

Redirecting metabolic flux to improve glucose paired glutamine/indigoidine formation is quantifiable across several other metrics. We should observe high TRY for our desired product since higher glutamine yields, to support growth, should result in more indigoidine yields. The production of indigoidine would shift from stationary phase to exponential phase, as the metabolism of glucose catabolism and glutamine production are paired. Finally, these phenotypes should maintain fidelity across a range of growth modes and scales.

We tested to ensure that indigoidine production was improved in the engineered strain relative to the controls in several laboratory cultivation formats. We tested production using both the native glucose and engineered galactose pathways as carbon sources. Both strains were cultivated with either 10 g/L glucose or galactose, as the same targeted reaction set would function on either carbon source. In a deep well plate format, we observed that the engineered strain produced nearly three-fold more indigoidine than the control strain when fed glucose (**Figure 3a**). In a shake flask format, the engineered strain produced 30% more than the control strain. Finally, when cells were cultivated with galactose in the deep well format, the same engineered strain was able to produce indigoidine in contrast to the galactose utilization control strain which only formed biomass (**Figure 3b**).

**Figure 3:**
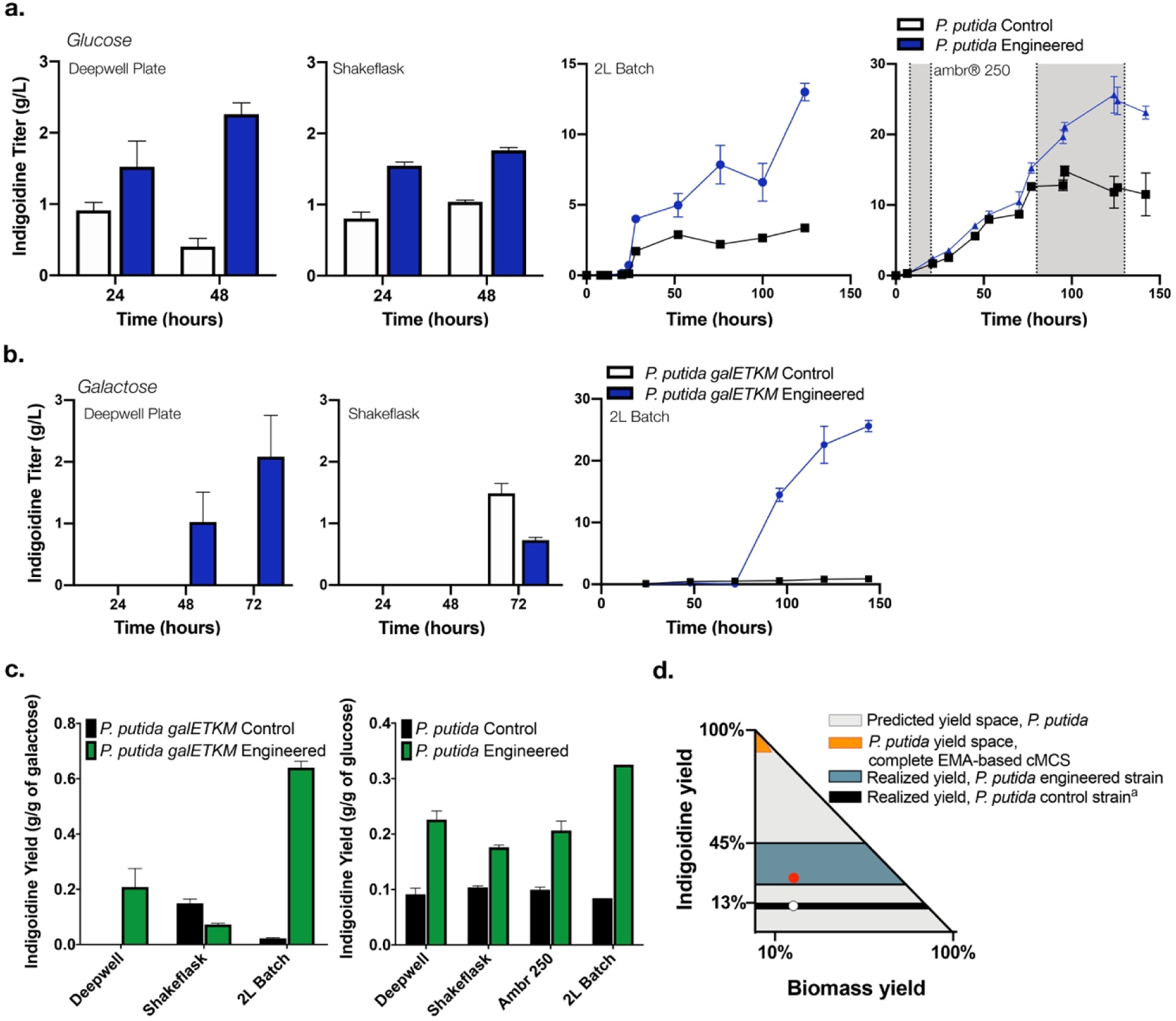
The product substrate pairing approach can improve Titer, Rate and Yield (TRY) across two carbon sources. A. Analysis of *P. putida galETKM* multi-gene engineered strains and a control strain (*P. putida galETKM*, empty vector plasmid) for production of indigoidine using glucose (**a**) or galactose (**b**) as the sole carbon source in M9 minimal medium. The culture format assessed is indicated above each panel. A fed-batch mode of cultivation was implemented in the ambr® 250 cultivation format. Glucose feeding is indicated by the gray shaded area. Control samples indicated with black outlined bars or black dots. The multi-gene engineered strains are indicated with blue bars or blue dots. **c**, Analysis of indigoidine yield across cultivation formats for both glucose-fed and galactose-fed strains. Yield from the control strain is indicated with black bars, and the multi-gene engineered strain is indicated with green bars. **d**, Predicted production envelope using genome scale model and constraint-based methods represented as theoretical yields of indigoidine as a function of biomass yields. Possible yield space for control strain is represented in grey. The possible yield space for 16 gene cMCS predicted by EMA is represented in orange. The range of observed experimental yield space for either the control or engineered strain across different production formats is represented with black and teal fill. A red dot indicates the realized production yield vs biomass yield in exponential phase under optimized conditions. The phase shift in production from stationary phase to exponential is not depicted.

In a 2L bioreactor, cultivated in a batch-mode with glucose as the carbon source, we observed improved titers at 12.5 g/L indigoidine from 60 g/L glucose. The control production strain, in contrast, produced 5 g/L, and production of the final molecule was realized after glucose was exhausted from the media. When galactose was fed, the engineered strain also exhibited improved titers of 25.6 g/L of indigoidine from 60 g/L galactose as opposed to the control strain that generated only around 900 mg/L of indigoidine; a 28-fold improvement in production was observed in the engineered strain. Moving to an industrially relevant cultivation format did not impact the final product titer, allowing us to further develop cultivation methods by switching to fed-batch mode.

We realized greater improvements in final product titer as well as improvements in production kinetics in the fed-batch mode using the ambr® 250 system. After administering an initial high nutrient feed to increase biomass in the reactor, we reduced the feed rate to study indigoidine product formation during exponential phase growth **(Figure 3a**, right hand panel, and **Supplementary Figure 3)**. During this phase, the engineered strain produced at a rate of 0.22 g/l/h, while the control strain accumulated no additional product. This observation is consistent with our hypothesis that indigoidine formation would occur during exponential phase due to pairing with glucose. In terms of yield, the engineered strain generated consistently higher production than the control strain when cultivated with glucose (0.2 g/g compared to 0.1 g/g), but was not as consistent when cultivated on galactose **(Figure 3c)**. Together all aspects of the phenotypes that were desirable for the engineered strain were found to be true.

## Discussion

This study is the first implementation of cMCS predictions enabled with CRISPR interference and resulted in a strain where production was paired with growth. Pairing the final desired product with the carbon source used for growth, mimics native obligatory product formations such as ethanol production and results in high productivity at scale. Further, to our knowledge, there are no other reports where the production of a non-native molecule was shifted from stationary phase to exponential phase as a result of strain engineering.

The competition between biomass accumulation and production of the target compound is a well-recognized challenge in biomanufacturing. This trade-off impacts both TRY and scalability. Approaches to address this tradeoff range from growth coupling^29^ to growth decoupling.^30^ Canonical approaches to growth coupling are FBA-based methods such as OptKnock^31^ that identifies secondary pathways that reduce the pool of a key intermediate as means to increase flux to the target of interest. This strategy has been termed as “weak” growth coupling^32^ where growth still occurs even if the desired product is not formed. Yim *et al.* used a tailored solution involving such computational methods to improve 1,3-BDO production to 18 g/L^33^, but their method cannot be generalized for other molecules. Others have described growth coupling as the creation of a “driving force” such as ATP production or cofactor imbalance, and link the driving force to the desired production pathway^29,34–36^. Driving force coupling is also pathway specific and requires additional strain engineering. Examples include 1-butanol production in *E. coli* using NADH as the driving force^34^ or media supplementation for butanone production in *E. coli* linked to acetate assimilation^29^.

In contrast to the examples described above, the multi-gene engineered product substrate pairing we report here is an implementation of “strong” growth coupling. It relies on EMA based methods that provide targets at the genome-scale level^11,37,38^ but predicts a large number of enzymatic reactions for elimination. We used FBA to corroborate our optimal cMCS and removed essential genes from targeted gene sets using -omics data to determine the genes that should be targeted for CRISPRi. This genome scale approach also represents a valuable paradigm for the evaluation of microbial hosts for their production capacity and could significantly reduce the time taken to optimize carbon source conversion to the final product. The appeal of this strategy is that the gene knockdown solution is scale-agnostic; the predicted metabolic rewiring should apply even in the largest bioreactor formats.

In the context of TRY improvement alone, indigoidine itself is an example of a heterologous product that has been demonstrated at high titers^39–41^. The production of indigoidine was high in the oleaginous yeast *Rhodosporidium toruloides* but remained low in the model yeast *S. cerevisiae*, despite similar optimization of cultivation parameters. This comparison represents an empirical example of the innate metabolic potential of a given host, and is consistent with our calculated max theoretical yields for indigoidine (**Supplementary Table 2**). Genome scale metabolic models can accurately predict how microbial hosts could be advantageous for the production of a given metabolite. For indigoidine, the MTY from glucose in *P. putida* is 0.54 mol/mol and is comparable to that for *R. toruloides* (0.5 mol/mol), while *E. coli* (0.4 mol/mol) and *S. cerevisiae* (0.079 mol/mol) are much lower. It is likely that every molecule will be different. Thus, selecting the best host/final product pair is a crucial aspect of developing the ideal production platform.

While our engineered strains showed many desirable phenotypes, several aspects merit additional discussion. The predicted EMA based cut set (cMCS) demands zero flux through these reactions for strong growth coupling. We excluded two genes from the predicted gene set due to their essentiality. Of the remaining gene targets, our RNAseq and proteomic data indicates a partial gene knockdown, implying that a non-zero flux could occur through the predicted reactions. The resulting yield space for indigoidine production is now different from what was predicted by EMA **(Figure 3d).** This suggests that partial implementation of the EMA predictions, still resulted in a shift of production from stationary to exponential phase while maintaining desirable indigoidine TRY.

Our approach allowed us to achieve, in one cycle of strain engineering, a high and consistent TRY for indigoidine across cultivation scales. With improvements in genetic tools and metabolic models it may be possible to obtain the 90% MTY predicted by the EMA based cMCS. A better understanding of the terminator sequence efficiency (as observed in this study and elsewhere in *E. coli*^*25*^) would enable more efficient CRISPR mediated gene knockdown. Similar fold repression of targeted proteins by CRISPRi/dCas9 were recently reported^42^, suggestive of a limitation for existing CRISPR systems in *P. putida*. Additionally, delineation of gene targets for this approach relies on the availability of high-quality genome scale metabolic models, and also calculated using a single carbon source. Future mechanistic studies of these strains will lead to more refined genome scale models, enabling more accurate metabolic flux modeling when the engineered strains are grown with these carbon sources. This approach cannot be used for certain mixed carbon streams, such as glucose and xylose, as our calculations for glucose pairing inactivates the pentose phosphate pathway. Similarly, there are metabolites that cannot be made obligatory for growth^11^. For final products derived from this class of metabolites, alternative strategies or hosts would need to be explored. We also do not take into consideration products or intermediates that may be toxic. As industrial processes also use renewable carbon sources that may contain inhibitory byproducts, microbial hosts will require some degree of tolerance engineering^43^ to unlock its potential. Addressing these aspects will further boost the usefulness of this product substrate pairing approach.

## Materials and Methods

### Computation of constrained minimal cut sets (cMCS)

*Pseudomonas putida* KT2440 genome scale metabolic model (GSM) iJN1462^9^ was used. The ATP maintenance demand and glucose uptake were 0.97 mmol ATP/gDW/h and 6.3 mmol glucose/gDW/h, respectively. Constrained minimal cut sets (cMCS) were calculated according to the algorithm as previously described^11^. Excretion of byproducts was initially set to zero, except for the reported overflow metabolites for secreted products specific to *P. putida* (gluconate, 2-ketogluconate, 3-oxoadipate, catechol, lactate, methanol, CO_2_, and acetate). Additional inputs including minimum demanded product yield (% of MTY) and minimum demanded biomass yield at 10% of maximum biomass yield were also specified in order to constrain the desired design space. Knockouts of export reactions and spontaneous reactions were not allowed. The algorithm computed for all minimal combinations of reaction knockouts blocking all undesired flux distributions and maintaining at least one of the desired metabolic flux distributions. With the specifications used herein each calculated knockout strategy (cMCS) will ensure that growth is not feasible without biosynthesis of glutamine. All cMCS calculations were done using API functions of CellNetAnalyzer^44^ on MATLAB 2017b platform using CPLEX 12.8 as the MILP solver. A summary of common potential growth coupled reactions and the number of targeted reactions to satisfy coupling restraints is included (**Supplementary Figure 4**). Once all the cMCS were enumerated, we used the decision workflow (**Figure 1a**) to identify an optimal engineering strategy for experimental validation.

### Constraint Based methods to confirm the cMCS

iJN1462 was extended to account for indigoidine biosynthesis pathway and checked for strong growth coupling to confirm the chosen engineering strategy for experimental implementation. Flux Balance Analysis (FBA) was used to calculate the maximum theoretical yield (MTY) from reaction stoichiometry and redox balance and also for single gene deletion analysis. Flux variability analysis (FVA) was used along with FBA to check for minimum and maximum glutamine or indigoidine flux under the identified cMCS strategy to confirm product obligatory growth. COBRA Toolbox v.3.0^45^ in MATLAB R2017b was used for FBA and FVA simulations with the GLPK (https://gnu.org/software/glpk), an open-source linear optimization solver. Production envelope was obtained using the internal COBRA Toolbox function, *productionEnvelope()* and plotted for *P. putida* (**Figure 3d**) as a fraction of maximum theoretical product yield on y-axis and maximum theoretical biomass yield on x-axis.

### Chemicals, media and culture conditions

All chemicals and reagents were purchased from Sigma-Aldrich (St. Louis, MO) unless mentioned otherwise. When cells were cultivated in a microtiter dish format, plates were sealed with a gas-permeable film (Sigma-Aldrich, St. Louis, MO). Tryptone and yeast extract were purchased from BD Biosciences (Franklin Lakes, NJ). Engineered strains were grown on M9 Minimal Media as described previously^46^ with slight modifications. Carbon sources (glucose or galactose) were used at 56mM and (NH_4_)_2_SO_4_ was used at 2 g/L, unless indicated otherwise.

### Strains and strain construction

*Pseudomonas putida* KT2440 was used as the host for strain engineering. All strains used in this study are described in **Supplementary Table 4** and are available upon request from public-registry.jbei.org. Upon publication, plasmid sequences are available at public-registry.jbei.org. Specific DNA sequences used to design the gRNA array are described in **Supplementary Dataset 1.** Electroporation with the respective plasmid was performed using a Bio-Rad (Bio-Rad Laboratories, Hercules, CA) MicroPulser preprogrammed EC2 setting (0.2 cm cuvettes with 100 µL cells, ∼5msec pulse and 2.5kV) with slight modifications^47^. Cells transformed with replicative plasmid DNA were allowed to recover at 25 °C for 2.5 hours before plating on selective agar media at 23 °C for overnight incubation. Cells transformed with non-replicative (integrating) plasmids were allowed to recover for 4-8 hours in LB media before plating on selective agar media at 23 °C for an additional 24 hours. Kanamycin sulfate or gentamicin sulfate (Sigma-Aldrich, St. Louis, MO) was used at a concentration of 50 µg/mL or 30 µg/mL, respectively. Integration of the *Ec.galETKM* operon or heterologous indigoidine gene pathway was implemented using a sucrose counterselectable plasmid for allelic exchange^48^. Positive clones were confirmed for the genotype by colony PCR using Q5 Polymerase enzyme (NEB, Ipswitch, MA). The dCpf1/CRISPRi system was adapted for use in *P. putida* by subcloning an endonuclease dead *Francisella tularensis subsp. Novicida Cpf1*^*49*^ into a pBBR1 backbone and placed under the LacUV5 promoter. The synthetic gRNA array was constructed using gene synthesis techniques (Genscript, Piscataway, NJ) and cloned into the dCpf1/CRISPRi backbone using isothermal DNA assembly. All plasmid constructs were verified with Sanger sequencing before transformation into *Pseudomonas putida*.

### Analytics/Sugar Quantification - HPLC

Glucose and organic acids from cell cultures were measured by an 1100 Series HPLC system equipped with a 1200 Series refractive index detector (RID) (Agilent Technologies, Santa Clara, CA) and Aminex HPX-87H ion-exclusion column (300 mm length, 7.8 mm internal diameter). 300 µL aliquots of cell cultures were removed at various time points during production and filtered through a spin-cartridge with a 0.45-μm nylon membrane, and 10 μL of the filtrate was eluted through the column at 50°C with 4 mM sulfuric acid at a flow rate of 600 μL/min for 30 min. Metabolites were quantified with an external standard calibration with authentic standards.

### Indigoidine Extraction and Quantification

Indigoidine was purified from *P. putida* with slight modifications as previously described^50^. Cells were lysed in 1% SDS and 100 mM NaCl and then centrifuged at 14,000 ×*g* for 3 minutes. The supernatant was discarded and the pellet was washed with three rounds of methanol, isopropanol, water, ethanol, and hexane to remove contaminating proteins and metabolites. The pellet was allowed to dry overnight and then resuspended in DMSO at a final concentration of 1mg/mL. Indigoidine purity was characterized by NMR. A standard curve correlating indigoidine concentration to OD_612_ was generated using this reagent (**Supplementary Figure 5**). The purity of extracted indigoidine (**Supplementary Figure 6**) from both *E. coli* and *P. putida* were cross-validated by NMR as previously described^41^.

To rapidly quantify indigoidine production in a high throughput manner, the colorimetric assay was used to determine indigoidine production. Briefly, pelleted 100uL of cells by centrifugation at 15000 rpm for 2 min. The supernatant was discarded and 500uL DMSO was added to the pellet. The solution was vortexed vigorously for 10 minutes to dissolve indigoidine. After centrifugation at 15000 rpm for 2 min, 100μL of DMSO extracted indigoidine was added to 96-well flat-bottomed microplates (BD Biosciences, San Jose CA). Indigoidine was quantified by measuring the optical density at using a microplate reader (Molecular Devices, San Jose, CA) preheated to 25°C and applying standard curve generated from indigoidine. The equation used was Y (g/L of Indigoidine) = 0.212*A_612_ - 0.0035. DMSO-solubilized cell lysate from wild-type *P. putida* does not contribute any signal when measured at OD_612_.

To correlate indigoidine yields with biomass yields, the dry cell weight was determined using OD_612_ to biomass conversion estimates as previously described^51^.

### RNAseq and data analyses

Total RNA was prepared following the manufacturer’s protocol^52^ for Trizol-based RNA extraction with several modifications. RNA from trizol treated lysates were bound to a silica column (Direct-zol RNA MiniPrep Plus Kit, Zymo Research, Irvine CA) and its integrity confirmed using a Bioanalyzer RNA 6000 Nano assay (Agilent Technologies, Santa Clara, CA). rRNA was removed from 100 ng of total RNA using Ribo-Zero(TM) rRNA Removal Kit (Illumina Biotechnology, San Diego, CA). Stranded cDNA libraries were generated using the Illumina Truseq Stranded mRNA Library Prep kit. The rRNA depleted RNA was fragmented and reversed transcribed using random hexamers and SSII (Invitrogen-ThermoFisher, Carlsbad, CA) followed by second strand synthesis. The fragmented cDNA was treated with end-pair, A-tailing, adapter ligation, and 10 cycles of PCR amplification. Prepared libraries were quantified using KAPA Biosystem’s next-generation sequencing library qPCR kit (Kapa Biosystems / Roche AG, Basel, Switzerland) and run on a Roche LightCycler 480 real-time PCR instrument. Sequencing of the flowcell was performed on the Illumina NovaSeq sequencer using NovaSeq XP V1 reagent kits, following a 2×150nt indexed run protocol. Reported gene expression values are the total normalized transcripts per million (TPM). All raw data is available through NCBI-SRA associated with NCBI-Bioproject (Accession IDs: PRJNA580539 - PRNJA580574) and the DOE-JGI IMG database (Project ID: 505977).

### Targeted proteomics by LC-MS/MS

A targeted SRM (selected reaction monitoring) method was developed to quantify relative levels of pathway proteins in samples under the various tested conditions in a 60 mL cultivation format. At the time points indicated, 1 mL of each sample was pelleted by centrifugation at 14,000*g* and flash frozen with liquid nitrogen at - 80 °C until ready for processing. Cells were lysed in 100 mM NaHCO_3_ using 0.1 mm glass beads using a Biospec Beadbeater (Biospec Products, Bartlesville, OK) with 60 s bursts at maximum power and repeated three times. Cell lysates were cooled on ice between each round. The clarified supernatant was harvested by centrifugation at 14,000*g*. The lysate protein concentration was estimated following the manufacturer’s directions for the BCA method (ThermoFisher Scientific/Pierce Biotechnology, Waltham, MA). The SRM-targeted proteomic assays and analyses were performed as described previously^53,54^. The SRM methods and data are available at Panoramaweb (shorturl.at/rsAK3).

### Cultivation at different scales

Cultures from glycerol stocks were struck to single colonies on LB agar media with the appropriate antibiotic as necessary. Single colonies were used to inoculate overnight cultures in LB with the appropriate antibiotic. Saturated overnight LB cultures were then back-diluted 1/100x into M9 minimal media with the appropriate carbon source as indicated. Cultures were back-diluted and adapted twice to ensure robust cell growth before heterologous pathway induction. Adaptation of *P. putida Ec.galETKM* strains for growth in M9 minimal salt media with galactose had a long initial adaptation phase of around 96-120 hours before cultures showed turbidity. All cultures were incubated with shaking at 200 rpm and 30°C. To prepare cells for pathway induction, M9 adapted cultures were back-diluted to a starting OD600 of 0.1, at which point IPTG and arabinose were added as appropriate. Production cultures grown in 24 well deep well plates (Axygen Scientific, Union City, CA) inoculated into a 200 µL culture volume and incubated InFors Multitron HT Double Stack Incubator Shaker (Infors HT, Bottmingen-Basel, Switzerland) set to 999 rpm linear shaker, and 70% humidity. For shake flask experiments, 60 mL cultures were grown in 250mL unbaffled Erlenmeyer shake flask and incubated at 200 rpm with orbital shaking. For all experiments, the indigoidine pathway was induced with 0.3% w/v L-arabinose, and dCpf1 mediated gene repression was induced with 500 µM IPTG. Production assays were performed in independent biological triplicate and repeated at least twice, except for the scale up experiments (described below), which were performed in biological duplicate.

### Batch experiments at 2 L bioreactor scale

Batch experiments were performed using a 2 L bioreactor equipped with a Sartorius BIOSTAT B® fermentation controller (Sartorius Stedim Biotech GmbH, Goettingen, Germany), fitted with two Rushton impellers fixed at an agitation speed of 800 rpm. Initial reactor volume was 1 L M9 Minimal Media (10g/L Glucose, 0.3% w/v L-arabinose, 30mM NH_4_^+^), and 50 mL overnight pre-culture in the same media. Feeding solution contained 100 g/L glucose, 300mM NH_4_^+^ along with L-arabinose and kanamycin. The temperature was held constant at 30°C. The bioreactor pH was monitored using the Hamilton EasyFerm Plus PHI VP 225 Pt100 (Hamilton Company, Reno, NV) and was maintained at a pH of 7 using 10 M sodium hydroxide. Dissolved oxygen concentration was monitored using Hamilton VisiFerm DO ECS 225 H0.

### Advanced micro bioreactor method: 250 mL ambr® 250 bioreactor cultivations

Fed-batch bioreactor experiments were carried out in a 12-way ambr® 250 bioreactor system equipped with 250 mL single-use, disposable bioreactors (microbial vessel type). The vessels were filled with 150 mL M9 minimal salt media containing 10 g/L glucose as carbon source. Temperature was maintained at 30 °C throughout the fermentation process and agitation was set constant to 1300 RPM. Airflow was set constant to 0.5 VVM based on the initial working volume and pH was maintained at 7.0 using 4 N NaOH. Reactors were inoculated manually with 5 mL of pre-culture cell suspension. After an initial batch phase of 12 hours, cultures were fed with a concentrated glucose feed solution (600 g/L glucose, 120 g/L ammonium sulfate, 50 µg/mL kanamycin, 3 g/L arabinose and 500 µM IPTG) by administering feed boluses every two hours restoring glucose concentrations to 10 g/L (feed parameters: 3.1 min @ 50 mL/h). After observing glucose accumulation, feed addition was paused and resumed at reduced feed rates when glucose levels dropped below 10 g/L (1 min @ 50 mL/h). Experiments with a continuous feeding regime were initially fed at 1.3 mL/h (0.3 mL/h after seeing glucose accumulation). Samples were taken 1-2 times every day (2 mL) and stored at −20 °C. The ambr® 250 runtime software and integrated liquid handler was used to execute all process steps unless stated otherwise.

## Acknowledgements

We thank Héctor García Martin, Morgan Price, and Adam Deutschbauer (LBNL) for technical assistance and helpful comments on this work. The genomic analysis was conducted at the U.S. Department of Energy Joint Genome Institute, a DOE Office of Science User Facility, supported by the Office of Science of the U.S. Department of Energy under Contract No. DE-AC02-05CH11231. The design and engineering of microbial strains was conducted at the DOE Joint BioEnergy Institute (http://www.jbei.org) supported by the US Department of Energy, Office of Science, through contract DE-AC02-05CH11231 between Lawrence Berkeley National Laboratory and the US Department of Energy. Demonstration of scale-up was conducted at the Advanced Biofuels and Bioproducts Demonstration Unit (ABPDU). The United States Government retains and the publisher, by accepting the article for publication, acknowledges that the United States Government retains a non-exclusive, paid-up, irrevocable, world-wide license to publish or reproduce the published form of this manuscript, or allow others to do so, for United States Government purposes. In accordance with the journal’s conflict of interest policy, AM, TE, and DB have submitted a patent related to some of the work in this study.

## Contributions

Conceptualization of the project: AM, TE, DB. Development and implementation of Computational Methods: DB. Strain construction, molecular biology, analytical chemistry: TE, AL, RH, BW. Contributed critical reagents: TE, YS, RH, CP. Proteomic analysis: YC, CJP. RNAseq pipeline: VRS. NMR analysis: AL, YL. Interpreted results: TE, AL, DB, YC, JDK, AM. Bioreactors and Scaleup: AL, JPP, TE, AL. Drafted the manuscript: DB, TE, CJP, JDK, AM. Raised funds: AM and JDK. All authors contributed to and provided feedback on the manuscript. All authors read and approved the final manuscript.

## Description of Supplemental Figures, Tables, and Datasets

**Supplementary Figure 1:** Characterization of Indigoidine Production Kinetics in *Pseudomonas putida*.

**Supplementary Figure 2:** Quantification of CRISPRi efficacy in *Pseudomonas putida*.

**Supplementary Figure 3.** Replicate fed-batch ambr250 cultivation (continuous feeding regime) of CRISPRi engineered product substrate paired indigoidine production strategy.

**Supplementary Figure 4.** Output from Computational Growth Coupling Metabolic Modeling.

**Supplementary Figure 5:** Characterization of Indigoidine.

**Supplementary Figure 6**: Analysis of indigoidine purity by H-NMR.

**Supplementary Table 1:** Potential for product substrate pairing for all metabolites in *Pseudomonas putida* KT2440 and *E. coli* MG1655 using glucose as the sole carbon source.

**Supplementary Table 2:** Comparison of industrially relevant hosts for glutamine and indigoidine production.

**Supplementary Table 3:** Analysis of suitable starting carbon sources to determine compatible carbon sources for cultivation for substrate-product pairing with indigoidine.

**Supplementary Table 4:** Strains Used in this Study.

**Supplementary Data Set 1**: Gene Sequences Used Design of Synthetic CRISPR Interference gRNA Array.

**Supplementary Data Set 2**: Identification of essential genes in *P. putida* using barcoded transposon mutagenesis (RB-TnSeq).

