## Supplemental Figures and Tables for "Genome-scale metabolic rewiring to achieve predictable titers rates and yield of a non-native product at scale"

**Supplementary Figure 1: Characterization of Indigoidine Production Kinetics in *Pseudomonas putida*.**

(a) Kinetic time course of indigoidine production in *P. putida* KT2440. 60mL Cells were cultivated in M9 minimal media under standard conditions. Samples were harvested at the time points indicated to monitor optical density, extracellular glucose, and indigoidine. Sample time points are indicated as hours after 0.3% (w/v) arabinose induction.

(b) Comparison of indigoidine production in *P. putida* *galETKM* cultivated in 1% glucose or 1% galactose in M9 minimal salt media.

(c-e) Comparison of Carbon/Nitrogen (C/N) ratios for glucose and ammonium sulfate. *P. putida KT2440* carrying an genomically integrated heterologous indigoidine production pathway was cultivated in M9 minimal salt media where the glucose or ammonium sulfate concentration was varied as indicated. Samples were harvested 24 h or 48 h post induction with 0.3% (w/v) arabinose and indigoidine titer was measured as described (materials and methods). In (c), the same C/N ratio was compared but at two different concentrations of glucose and ammonium sulfate.

(f) Evaluation of native tRNA promoters. One hundred (100) bp promoter sequences from *P. putida* KT2440 tRNA ligases immediately upstream of the start ATG were amplified and cloned upstream of a *RBS-mCherry* gene cassette. Constructs were transformed into KT2440 grown in M9 minimal media three rounds of adaptation. mCherry signal was determined by flow cytometry after 24 hours of growth in a 24 well deep well plate. mCherry fluorescence was compared against background fluorescence in a control strain harboring an empty vector control (pTE219).

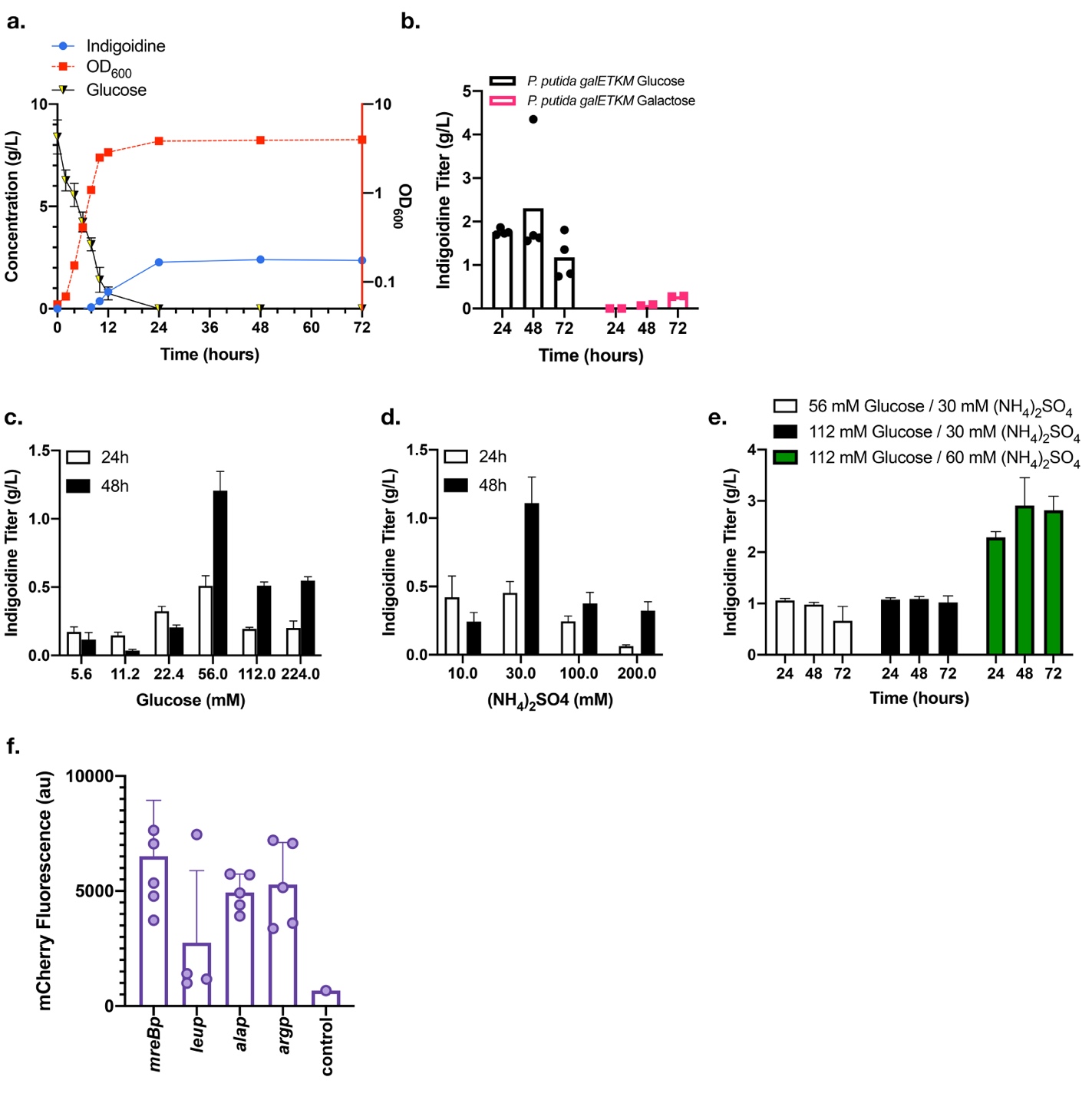

**Supplementary Figure 2: Quantification of CRISPRi efficacy in *Pseudomonas putida*.** RNAseq (A) and Proteomic (B) validation of dCpf1 multiplex targeted gene knockdown. *P. putida* strains harboring a genomically integrated indigoidine pathway and a plasmid-borne CRISPRi/dCpf1 engineered system were prepared for the production of indigoidine (Materials and Methods) and sampled at the indicated time points. No peptides from PP_4947p were detected (n.d.).

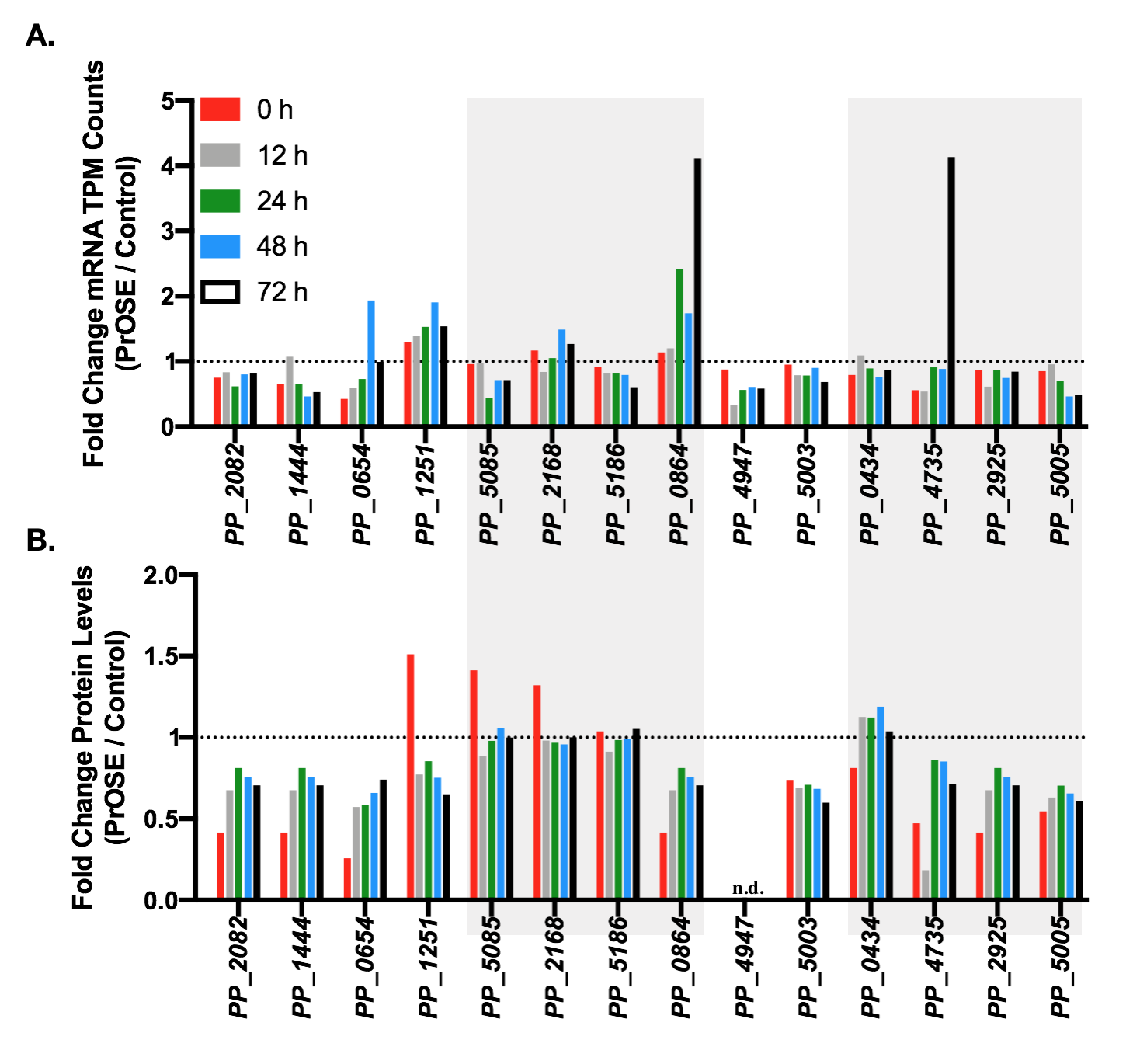

**Supplementary Figure 3. Replicate fed-batch ambr250 cultivation (continuous feeding regime) of CRISPRi engineered product substrate paired indigoidine production strategy.** Similar to the results described in Figure 3A, a replicate fed-batch feeding regime was implemented to demonstrate production of indigoidine during exponential phase growth. Instead of a bolus feeding regime as used in 3A, this replicate tested a continuous feeding regime. A similar production of indigoidine during feeding was observed (second gray area on graph) when the control strain did not produce additional indigoidine.

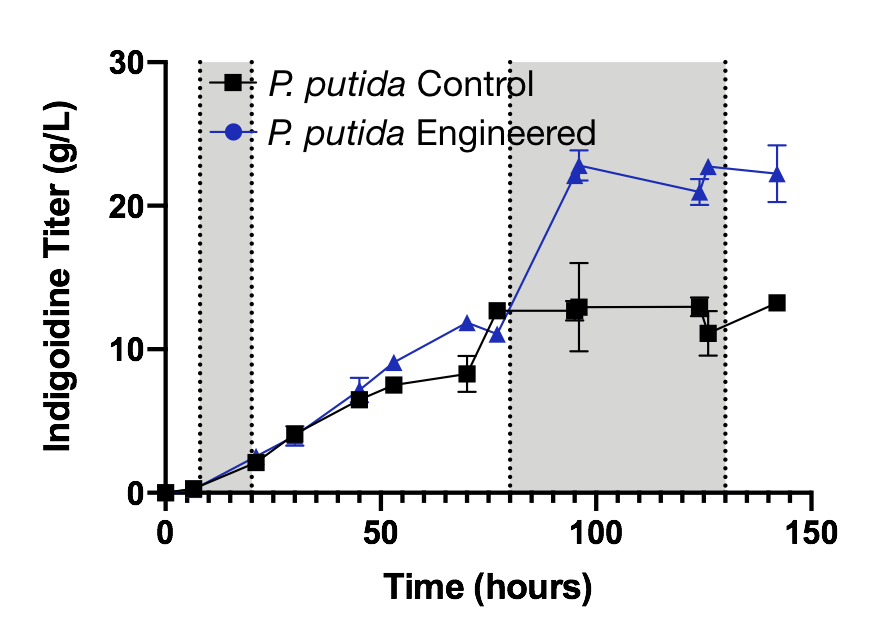

**Supplementary Figure 4. Output from Computational Growth Coupling Metabolic Modeling.** EMA based cMCS predictions for 417 metabolites that could be coupled to at least 10% biomass yield and a minimum production yield of 10%, 50% and 80% MTY. Each metabolite is represented as a triangle and the number of cut sets is shown in blue whereas size of cut set is shown in pink. The black line marks the median for each case.

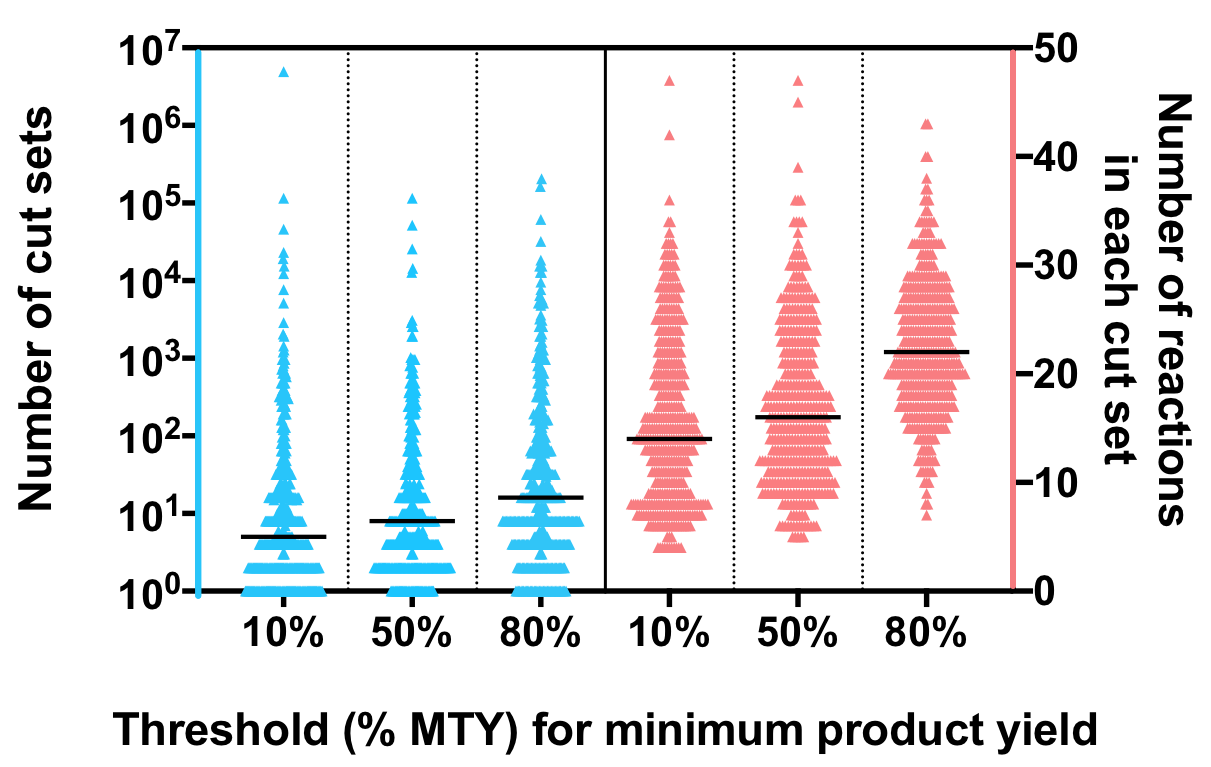

**Supplementary Figure 5: Characterization of Indigoidine.** A standard curve used for indigoidine quantification relating absorbance at wavelength 612 nm to indigoidine concentration dissolved in DMSO. In each equation Y is indigoidine concentration in g/L and x is absorbance at 612 nm (Materials and methods). The plotted standard curve for indigoidine purified from *P. putida* was repeated >3 times over from freshly generated microbial cultures over the course of several months. A standard curve was also generated using the same pathway genes expressed in *E coli*.

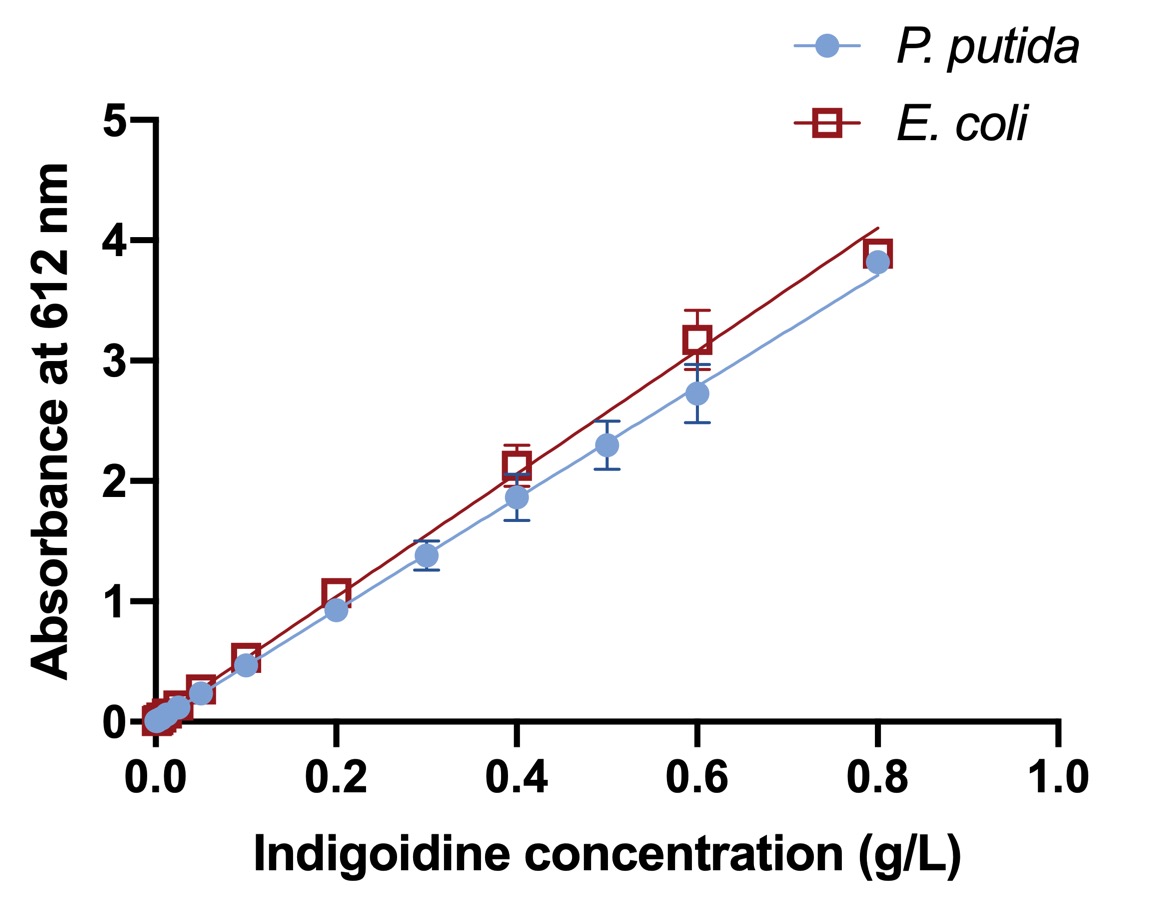

| Strains | Equation | R^2^ |
| --- | --- | --- |
| *P. putida* | Y = 0.212x - 0.0035 | 0.9911 |
| *E. coli* | Y = 0.196x - 0.0035 | 0.9921 |

**Supplementary Figure 6**: Analysis of indigoidine purity by H-NMR. Indigoidine was extracted from *P. putida* and *E. coli* harboring expression the heterologous pathway and purified as described (Materials and Methods). Indigoidine purified from either host shows similar purity.

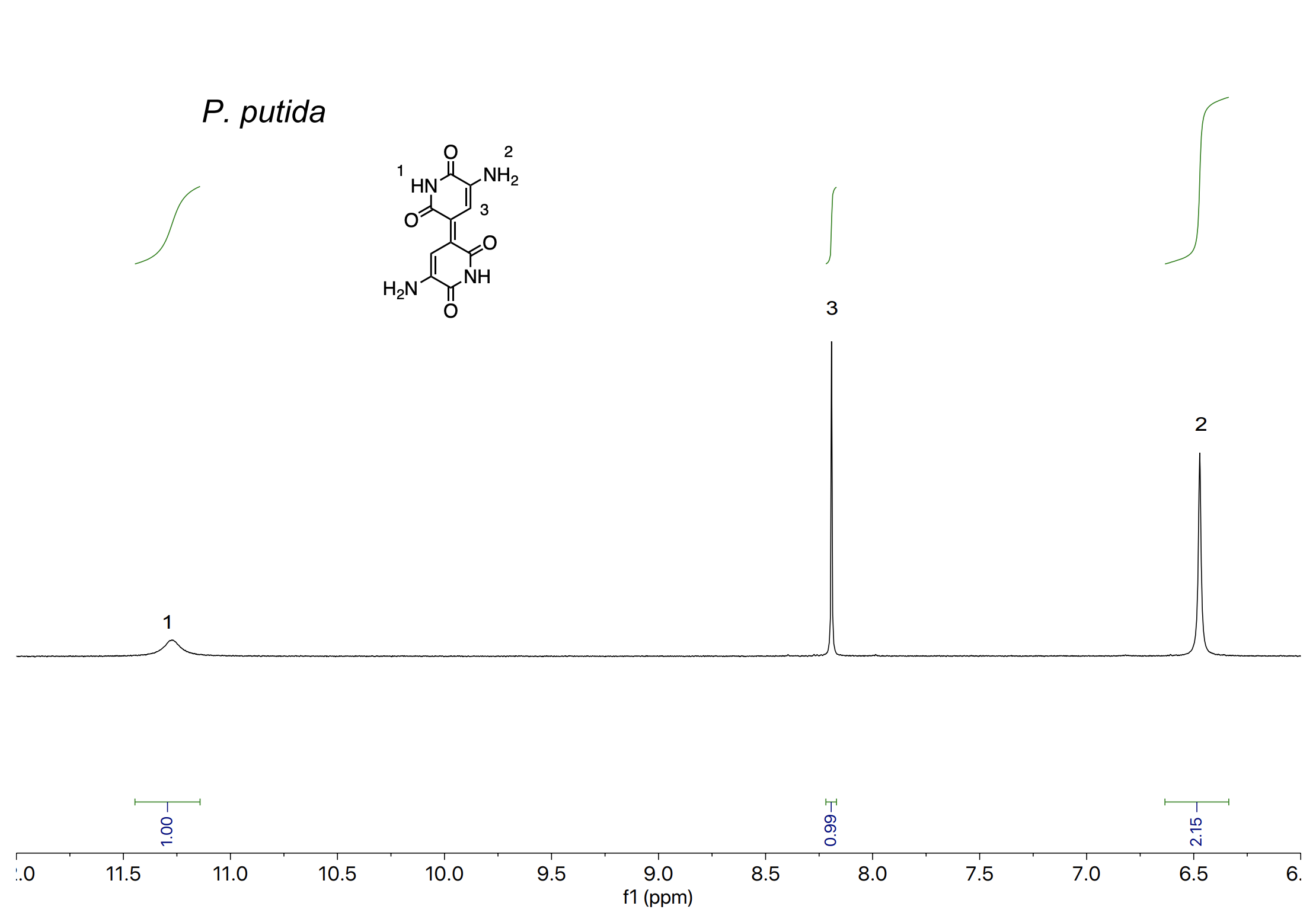

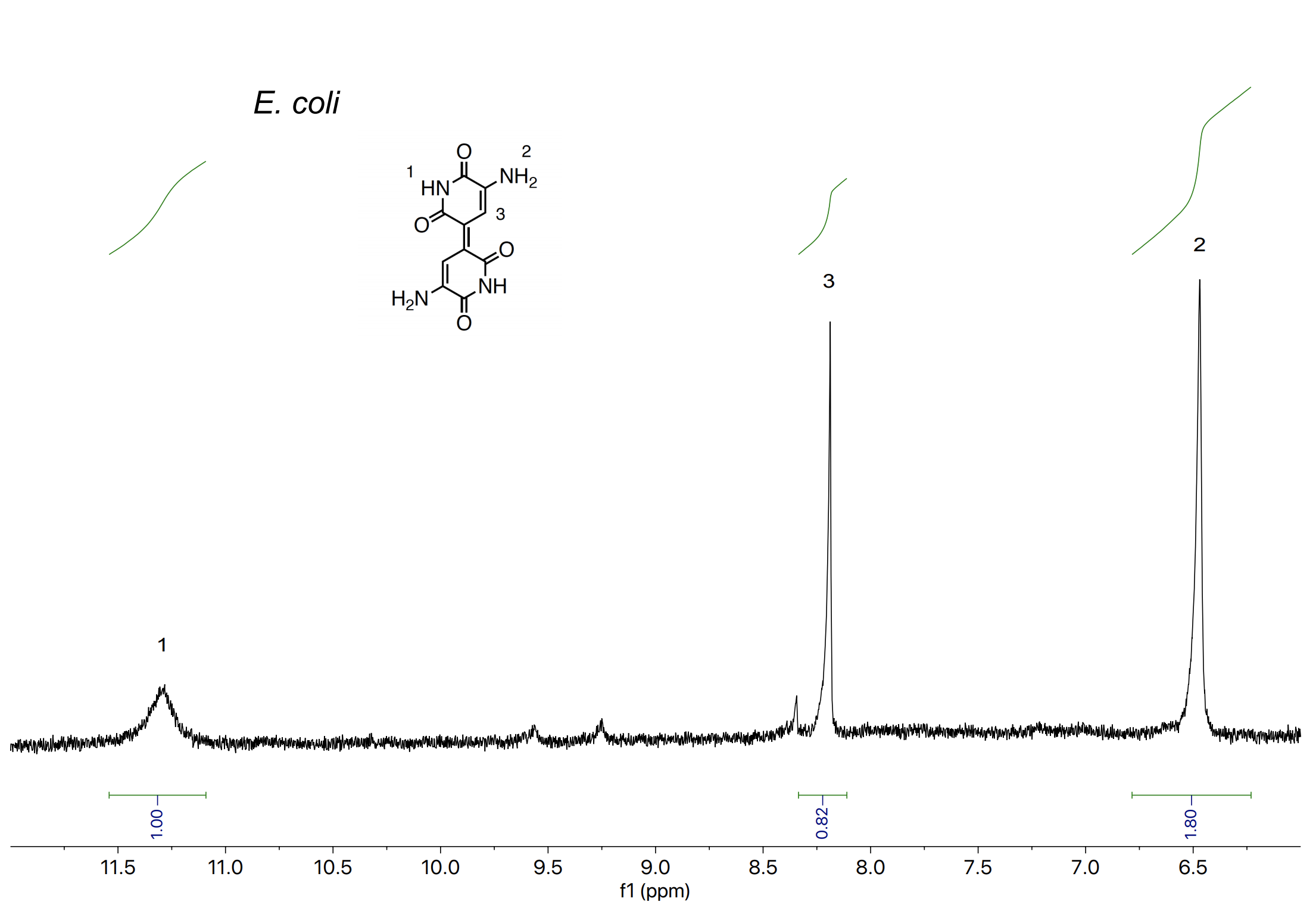

**Supplementary Table 1:** Potential for product substrate pairing for all metabolites in *Pseudomonas putida* KT2440 and *E. coli* MG1655 using glucose as the sole carbon source.

| **Organism** | | ***E. coli* MG1655**^1^ | ***P. putida* KT2440** | |
| --- | --- | --- | --- | --- |
| **Genome-scale model (GSM)** | | **iJO1366** | **iJN1462** | **Remarks** |
| **Specific constraints** | Glucose uptake limit (mmol/gDW/h) | 15 | 6.3 |  |
|  | ATP maintenance (mmol/gDW/h) | 3.15 | 0.92 |  |
| **Reactions** | Total Reactions | 2582 | 2928 |  |
|  | Repressible reactions | 1414 | 2046 |  |
|  | Irrepressible reactions | 1168 | 882 |  |
| **Metabolites** | Internal metabolites | 1805 | 2145 | 892 metabolites present in *P. putida*, but absent in *E. coli* |
|  | Glucose-producible organic metabolites | 954 | 979 |  |
|  | Minimum yield (%) | 10 | 10 |  |
|  | Metabolites with feasibility of strong coupling | 954 | 966 | **98.6%** |
| **cMCS size** | Min | 3 | 4 | Metabolites including 4-Hydroxy-L-threonine, Mannose-6-phosphate, Mannuronate |
|  | Max | 50 | 48 | Vaccenyl coenzyme A |
|  | Mean | 17.4 | 18.9 |  |

**Supplementary Table 2:** Comparison of industrially relevant hosts for glutamine and indigoidine production.

| **Organism** | ***P. putida*** | ***C. glutamicum*** | ***E.coli*** | ***R. toruloides*** | ***S. cerevisiae*** |
| --- | --- | --- | --- | --- | --- |
| **Genome-scale model (GSM)** | iJN1462[^1^](http://f1000.com/work/citation?ids=7711227&pre=&suf=&sa=0) | iCW773[^2^](http://f1000.com/work/citation?ids=7313437&pre=&suf=&sa=0) | iML1515[^3^](http://f1000.com/work/citation?ids=4418035&pre=&suf=&sa=0) | iRhto1108C[^4^](http://f1000.com/work/citation?ids=7785128&pre=&suf=&sa=0) | iMM904[^5^](http://f1000.com/work/citation?ids=2038165&pre=&suf=&sa=0) |
| **Maximum theoretical yields** | | | | | |
| **Glutamine (mol/mol glucose)** | 1.141 | 1 | 1.135 | 1.118 | 0.481 |
| **Biomass (gDW/mmol)** | 0.098 | 0.092 | 0.088 | 0.075 | 0.029 |
| **Indigoidine (mol/mol glucose)** | 0.537 | 0.4* | 0.4 | 0.503* | 0.079* |
| **Constraints used for each GSM to best represent cell phenotype** | | | | | |
| **Glucose uptake rate (mmol/gDCW/hr)** | 6.3 | 4.67 | 10 | 5 | 10 |
| **ATP maintenance demand (mmol/gDCW/hr)** | 0.92 | 0 | 6.86 | 1.012 | 1 |
| **Highest experimentally reported indigoidine titers (g/L), rates (g/L/h) and yields (g/g)** | | | | | |
| **Titers (g/L)** | 25.6 from glucose minimal media  (this study) | n.d. | 7.08 from rich media[^6^](http://f1000.com/work/citation?ids=4915410&pre=&suf=&sa=0) | 18.04 from rich media[^7,8^](http://f1000.com/work/citation?ids=7026959,7692385&pre=&pre=&suf=&suf=&sa=0,0) | 0.98 from rich media[^9^](http://f1000.com/work/citation?ids=6146654&pre=&suf=&sa=0) |
| **Rate (g/L/h)** | 0.22^a^ |  |  | 0.15 |  |
| **Yields  (g/g)** | 0.33^b^ |  |  | 0.19[^7,8^](http://f1000.com/work/citation?ids=7026959,7692385&pre=&pre=&suf=&suf=&sa=0,0) |  |

*No indigoidine formation if FMN is limiting.

^a^ When cells are fed glucose from a continuous feeding regime (i.e., exponential growth phase).

^b^ When cells are fed galactose in batch mode culture. (i.e., stationary phase)

**Supplementary Table 3:** Analysis of suitable starting carbon sources to determine compatible carbon sources for cultivation for substrate-product pairing with indigoidine.

| **Glucose/Indigoidine PrOSE modeling: acceptable alternate carbon sources** | **Glucose/Indigoidine PrOSE modeling: unacceptable alternate carbon sources** |
| --- | --- |
| Glycerol, Fructose, Mannose, Serine, Threonine, Galactose, Asparagine, Aspartate, Glycine, Homoserine, Xanthine | Lysine, Leucine, Succinate, Malate, Tryptophan, Tyrosine, Valine, Xylose, Proline, *p*-coumarate, Cysteine, Alpha-ketoglutarate |

**Supplementary Table 4:** Strains Used in this Study**.**

| **Strain Number** | **Genotype** | **Relevant Figure(s)** | **Reference** |
| --- | --- | --- | --- |
| JBEI-13809 | *Pseudomonas putida* KT2440 wild type prototroph CmR AmpR |  | ATCC 47054 Nieto *et al*[*^10^*](http://f1000.com/work/citation?ids=8197933&pre=&suf=&sa=0) |
| JBEI-137184^a^ (TEAM-1120) | KT2440 *Pp_5402::arap-Sc.bpsA,Bc.sfp* | Supplemental Figure 1 | This study |
| JBEI-137183^a^ (TEAM-1411) | KT2440 *Pp_5402::arap-Sc.bpsA,Bc.sfp Pp_0871::galp-Ec.galETKM* | Supplemental Figure 1 | This study |
| JBEI-137178 (TEAM-1361) | KT2440 {p/pTE328 *neo^d^ BBR1 PP_0933(mreB)p-RBS-mCherry*} | Supplemental Figure 1 | This study |
| JBEI-137179 (TEAM-1362) | KT2440 {p/pTE329 *neo BBR1 PP_4794(leu)p-RBS-mCherry*} | Supplemental Figure 1 | This study |
| JBEI-137180 (TEAM-1363) | KT2440 {p/pTE330 *neo BBR1 PP_4474(ala)p-RBS-mCherry*} | Supplemental Figure 1 | This study |
| JBEI-137181 (TEAM-1364) | KT2440 {p/pTE330 *neo BBR1 PP_5089(arg)p-RBS-mCherry*} | Supplemental Figure 1 | This study |
| JBEI-136298^b^ | KT2440 *Pp_5402::arap-Sc.bpsA,Bc.sfp* {p/pTE219 *neo BBR1 Placuv5-dCpf1* *j23101p-nt_gRNA*} | Figure 2, Figure 3, Supplemental Figure 2 | This study |
| JBEI-105555^c^ | KT2440 *Pp_5402::arap-Sc.bpsA,Bc.sfp* {p/pTE327 *neo BBR1 lacuv5p-dCpf1* 14 gene CRISPRi array} | Figure 2, Figure 3, Supplemental Figure 2 | This study |
| JBEI-136186^c^  (TEAM-1425) | KT2440 *Pp_5402::arap-Sc.bpsA,Bc.sfp Pp_0871::galp-Ec.galETKM*  {p/pTE219 *neo BBR1 Placuv5-dCpf1* *j23101p-nt_gRNA*} | Figure 2, Figure 3 | This study |
| JBEI-136187^b^ (TEAM-1423) | KT2440 *Pp_5402::arap-Sc.bpsA,Bc.sfp Pp_0871::galp-Ec.galETKM*  {p/pTE327 *neo BBR1 lacuv5p-dCpf1* 14 gene CRISPRi array} | Figure 2, Figure 3 | This study |
| JBEI-18378 | E. coli BL21(DE3) Δ*prp*RBCD::t7p-*sfp*,t7p-*prpE* | Supplemental Figure 4 | Pfeifer *et al*[*^11^*](http://f1000.com/work/citation?ids=686035&pre=&suf=&sa=0) and Wehrs *et al^9^* |

^a^ Genomic integrations are targeted to an intergenic region adjacent to the indicated locus.

^b^ nt: non-targeting gRNA sequence with no homology to the *P. putida* KT2440 genome.

^c^ Refer to Supplemental Table 4 for sequences of targeted genes with sequences for targeting gRNAs, promoters, and terminators used in the multiplex array.

^d^ *neo* is also known as APH(3')-II family aminoglycoside O-phosphotransferase and confers resistance to 50 µg/mL kanamycin in P. putida.

**Supplementary Data Set 1**: Gene Sequences Used Design of Synthetic CRISPR Interference gRNA Array.

**Supplementary Data Set 2**: Identification of essential genes in *P. putida* using barcoded transposon mutagenesis (RB-TnSeq).

*Both supplemental datasheets are included as an excel file:*

Banerjee and Eng et al Supplementary Datasets.xlsx
